# IRE1α/XBP1 pathway expression is impaired in pediatric cholestatic liver disease explants

**DOI:** 10.1101/2022.03.11.484034

**Authors:** Alyssa Kriegermeier, Angela Hyon, Brian LeCuyer, Susan Hubchak, Xiaoying Liu, Richard M. Green

## Abstract

**Background/Aims:** Cholestatic liver diseases (CLD) are the leading indication for pediatric liver transplantation. Increased intrahepatic bile acid concentrations cause endoplasmic reticulum (ER) stress and the unfolded protein response (UPR) is activated to maintain homeostasis. UPR dysregulation, including the inositol-requiring enzyme 1α/X-box protein 1 (IRE1α/XBP1) pathway, is associated with several adult liver diseases. We evaluated hepatic UPR expression in pediatric patients with end-stage CLD and hypothesize that an inability to appropriately activate the hepatic IRE1α/XBP1 pathway is associated with the pathogenesis of CLD.

**Methods:** We evaluated 34 human liver explants. Cohorts included: pediatric CLD (Alagille, ALGS, and progressive familial intrahepatic cholestasis, PFIC), pediatric non-cholestatic liver disease controls (autoimmune hepatitis, AIH), adult CLD, and normal controls. We performed RNA-seq, quantitative PCR, and western blotting to measure expression differences of the hepatic UPR and other signaling pathways.

**Results:** Metascape pathway analysis demonstrated that the KEGG ‘protein processing in ER’ pathway was downregulated in pediatric CLD compared to normal controls. Pediatric CLD had decreased hepatic IRE1α/XBP1 pathway gene expression and decreased protein expression of p-IRE1α compared to normal controls. These CLD changes were not disease-specific to ALGS or PFIC. IRE1α/XBP1 pathway gene expression was decreased in pediatric CLD compared to AIH disease controls.

**Conclusion:** Pediatric CLD explants have decreased gene and protein expression of the protective IRE1α/XBP1 pathway and down-regulated KEGG protein processing in the ER pathways. IRE1α/XBP1 pathway expression differences occur when compared to both normal and non-cholestatic disease controls. Attenuated expression of the IRE1α/XBP1 pathway is associated with cholestatic diseases and could be targeted to treat pediatric CLD.

## Introduction

Cholestatic liver diseases (CLD), including genetic diseases such as Alagille syndrome (ALGS) and progressive familial intrahepatic cholestasis (PFIC), are the leading indication for pediatric liver transplantation^1^. During cholestatic conditions, bile acids accumulate within the liver which can lead to cell death. Though the genetic mutations associated with many cholestatic diseases are known, a spectrum of severity exists and the underlying factors that influence disease progression are not fully understood. Standard medical therapy for pediatric cholestatic liver diseases is supportive care and there are currently no medical treatments that prevents progression to end-stage liver disease and need for liver transplantation.

A major factor responsible for the lack of effective treatments for pediatric CLD is the gap in our understanding of disease modifying factors, including the protective hepatic responses required to reduce bile acid-induced injury. Cholestasis and the accumulation of hepatic bile acids increases endoplasmic reticulum (ER) stress in animal models and bile acids alter protein folding in vitro^2–5^. ER stress is caused by an excess of unfolded or misfolded proteins in the ER and has been shown to be important in the pathogenesis of several liver diseases including NAFLD/NASH, chronic viral hepatitis, alpha-1-antitrypsin deficiency, and alcohol-related liver diseases^6–9^. The unfolded protein response (UPR) is a protective molecular response present in all eukaryotic cells that decreases ER stress within cells and restores normal cellular homeostasis. There are 3 pathways of the UPR: the inositol-requiring enzyme 1α/X-box protein 1 (IRE1α/XBP1) pathway, the PKR-like ER kinase (PERK) pathway and the activating transcription factor 6 (ATF6) pathway. The IRE1α/XBP1 pathway is the most evolutionarily conserved. In the presence of ER stress, activated phosphorylated IRE1α (p-IRE1α) splices XBP1 mRNA to produce the transcriptionally active spliced form XBP1s, although XBP1 independent pathways also exist. Though primarily protective, if ER stress is prolonged and unresolved, the UPR can trigger apoptosis^10^.

Our previous animal studies have demonstrated that UPR pathways regulate liver bile acid synthesis and transporters, and hepatocyte-specific deletion of the UPR protein XBP1 alters bile acid metabolism and susceptibility to liver injury^11–13^. Mice lacking hepatocyte XBP1 have normal activation of the UPR but have impaired ER stress resolution with resultant increased hepatic apoptosis^14^. We have demonstrated in murine models that young mice are more susceptible to bile acid-induced liver injury and age-dependent changes in IRE1α/XBP1 pathway expression may be an important factor in resolving the ER stress and liver injury^15^. However, studies of the hepatic UPR in human pediatric liver diseases have not been previously performed. Therefore, in this study, we sought to evaluate the hepatic UPR expression in the livers of pediatric patients with severe cholestatic disease. Based on our prior murine data, we hypothesized that there is inadequate hepatic IRE1α/XBP1 activation in these patients in response to ER stress which would contribute to increased hepatocellular injury and fibrosis. By identifying liver UPR pathways that are impaired in these conditions, novel, targeted treatment strategies can be developed for a spectrum of pediatric cholestatic liver diseases.

## Materials and Methods

### Materials

Antibodies are listed in Supplemental Table 1. PCR primers were synthesized by either Sigma (St. Louis, MO) or Integrated DNA technologies (Coralville, Iowa), and are listed in Supplemental Table 2. All other reagents were high grade and purchased from commercial vendors who performed QC.

### Human Samples

A total of 34 human livers were obtained from biorepositories at Ann & Robert H. Lurie Children’s Hospital (LCH), Chicago, Illinois. This study was approved by the LCH institutional review board. Liver tissue was collected in accordance with existing IRB regulations at the time of collection and included both pediatric and adult explants (from the adult transplant center) that were stored at LCH biorepositories. All diseased liver samples, including CLD and autoimmune hepatitis (AIH) samples, were explants from patients undergoing liver transplantation. Normal control livers were explants from deceased persons who had donated their liver for transplantation but were unable to be utilized for transplantation due to recipient issues. Explanted livers were frozen at −80°C. Samples included 9 pediatric (<18 years of age) normal (pNormal) livers, 10 pediatric CLD livers (pCLD), 5 pediatric AIH livers, 3 adult CLD livers (aCLD), and 7 adult (>18 years of age) normal (aNormal) livers. Based on prior known clinical and histopathologic diagnoses, the aCLD cohort was composed of 2 patients with primary biliary cholangitis (PBC) and 1 patient with primary sclerosing cholangitis (PSC), and the pCLD cohort was composed of 5 patients with ALGS and 5 patients with PFIC.

### RNA Extraction, Quantitative Real-time PCR

Total RNA was extracted from frozen liver samples using TRIZOL reagent (Ambion by Life Technologies, Carlsbad, CA), and cDNA was synthesized with the qScript cDNA synthesis kit (Quantabio, Beverly, MD). Quantitative PCR (qPCR) was performed using Power SYBR Green PCR Master Mix (Thermo Fisher Scientific, Waltham, MA) as previously described^12^. Relative expression of the genes of interest were measured by the ΔΔCt method using beta-2 microglobulin (B2M) as a reference gene.

### RNA-seq preparation and analysis

RNA for RNA-seq was submitted for all 34 samples (10 pCLD, 3 aCLD, 5 AIH, 9 pNormal, and 7 aNormal). The stranded mRNA-seq was conducted in the Northwestern University NUSeq Core Facility. Briefly, total RNA examples were checked for quality using RINs generated from Agilent Bioanalyzer 2100. RNA quantity was determined with Qubit fluorometer. The Illumina Stranded mRNA Library Preparation Kit was used to prepare sequencing libraries from 100 ng of high-quality RNA samples (RIN>7). All 34 samples met QC standards and were used for the study. The kit procedure was performed without modifications. This procedure includes mRNA purification and fragmentation, cDNA synthesis, 3’ end adenylation, Illumina adapter ligation, library PCR amplification and validation. lllumina HiSeq sequencer was used to sequence the libraries with the production of single-end, 50 bp reads at the depth of 20-25 M reads per sample. The quality of reads, in FASTQ format, was evaluated using FastQC. Reads were trimmed to remove Illumina adapters from the 3’ ends using cutadapt^16^. Trimmed reads were aligned to the human genome (hg38) using STAR^17^. Read counts for each gene were calculated using htseq-count in conjunction with a gene annotation file for hg38 obtained from Ensembl (http://useast.ensembl.org/index.html)^18^. Normalization and differential expression were calculated using DESeq2 that employs the Wald test^19^. The cutoff for determining significantly differentially expressed genes was an FDR-adjusted p-value less than 0.05 using the Benjamini-Hochberg method^20^. Pathway analysis was performed using Metascape to identify pathways enriched in the significantly differentially expressed genes^21^. Pathway analysis was conducted both by utilizing a maximum of 3000 genes based on most significant adjusted p-values, and by utilizing differentially expressed upregulated or downregulated genes based on log2 fold change. Heatmaps of differentially expressed genes were generated in R utilizing results of the DESeq2 analysis.

### Western Blotting

Protein homogenates from frozen liver tissue were isolated using T-PER protein extraction reagent (ThermoFisher Scientific, Waltham, MA) containing protease inhibitor cocktails and Halt phosphatase inhibitors (EMD Millipore, Billerica, MA). Protein quantification was performed using Coomassie Plus protein assay reagent (ThermoFisher Scientific, Waltham, MA). Equal amounts of protein samples were subjected to immunoblotting for target proteins and immunoreactive bands were visualized using the Amersham ECL western blotting detection kit according to the manufacturer’s protocol (GE Healthcare, Piscataway, NJ). Antibodies for GAPDH or Cyclophilin B were used as loading controls.

### Tissue Histology Analysis

Sections of frozen liver tissue were thawed overnight in formalin at room temperature, and histology including H&E and Massons-Trichrome staining was prepared by the Pathology Core Facility at Robert H. Lurie Comprehensive Cancer Center of Northwestern University. Two blinded investigators (AK, RG) scored trichrome histology slides for degree of fibrosis (F0-F4) based on METAVIR scoring by reviewing 10 fields of view at 10× magnification^22^.

### Statistics

Data was analyzed using Prism Software and shown as mean ± SEM for all experiments. Comparison between two groups was performed using two-tailed student’s t-tests, comparison between 3 or more groups was performed using ANOVA with Tukey’s multiple comparisons, and significance was defined as a p≤0.05.

## Results

### Demographic and clinical information

A summary of available demographic/clinical data for the 34 liver samples used in this study is listed in Supplemental Figure 1. The age and sex was available for all normal samples except for 1 pediatric sample. Pediatric normal samples ranged from 2 - 5 years (mean 3.4 years), and 3 out of 9 (33%) were male. The adult normal samples ranged from 19 - 66 years (mean 31.6 years) and 2 out of 7 (29%) were male. The patient age was available for 2 of 3 aCLD samples (49 and 62 years), and 1 of 3 samples was identified as female while the other 2 samples had no sex identified. For the pCLD samples, liver disease diagnosis was specified but the age and sex were not available although LCH does not routinely perform liver transplantation for patients over the age of 18. The mean AIH patient age was 11.2 years, range (1 – 14 years) and 1 of 5 (20%) AIH samples was male. The average direct bilirubin on the day prior to transplantation for the AIH cohort was 2.36 mg/dL (range <0.2-8.9 mg/dL). Notably, within the AIH cohort, only patients 1 and 5 had direct bilirubin levels >1mg/dL and these patients were transplanted during their initial disease presentation and met criteria for liver failure (INR>2) (Supplemental Fig.1B).

### All CLD and AIH liver samples show significant fibrosis

Thirty-two of 34 (94%) available samples had adequate sample size to grade for fibrosis. Pediatric CLD and AIH explant liver samples had similar degrees of fibrosis (3.4 ± 0.2 and 3.9 ± 0.1 respectively), but both groups demonstrated significantly more fibrosis than pNormal samples (0.7 ± 0.2, p<0.001), and aCLD samples had significantly more fibrosis than aNormal samples (4.0 ± 0.0 vs 0.6 ± 0.3, p<0.001) (Supplemental Fig. 2A, B). When ALGS and PFIC sub-groups were compared to AIH, both sub-groups had similar fibrosis (3 ± 0.4, and 3.7 ± 0.3, p=0.3), and both disease groups had more fibrosis than normal controls (p<0.001) but were similar to the AIH cohort (Supplementary Fig. 2C). This confirmed that all disease cohorts in this analysis were from patients with end-stage chronic liver disease.

### RNA-seq, principal component analysis (PCA), and pathway analysis of pCLD and to pNormal livers

Principal component analysis (PCA) of RNA-seq data demonstrated that all disease samples (CLD and AIH) cluster distinctly from normal pediatric and normal adult controls (Fig. 1A). Comparing pCLD to pNormal samples, there were 5,766 significantly differentially expressed genes (adj p-value <0.05) identified using RNA-seq. Metascape pathway analysis of the top 3000 genes with the lowest adjusted p-values demonstrated that genes involving KEGG pathway hsa04141 “*protein processing in endoplasmic reticulum*” was among the top 5 differentially expressed pathways (−log10(P)=25.77) (Fig. 1B). Heatmap analysis demonstrated that expression of most genes within this pathway were decreased in the pCLD samples compared to pNormal livers (Fig. 1C). To confirm that this pathway was significantly downregulated in pCLD, Metascape pathway analysis of all downregulated genes (n=2707) in pCLD compared to pNormal was performed and again identified this same KEGG pathway as one of the top 10 significantly differentially expressed pathways (−log10(P)=35.69) (Supplemental Fig. 3A). In contrast, utilizing the top significantly upregulated genes in the pCLD samples, this pathway was not within the top 100 differentially expressed pathways (Supplemental Fig. 3B). We next performed a sub-group analysis of the pCLD cohort and compared PFIC to ALGS samples. RNA-seq identified 351 significantly differentially expressed genes between PFIC and ALGS samples, although pathway analysis did not demonstrate that the KEGG pathway hsa0414 was differentially expressed between these two cholestatic liver diseases, nor were other pathways involved in protein folding or ER stress (Supplemental Fig. 4A).

**Figure 1.**
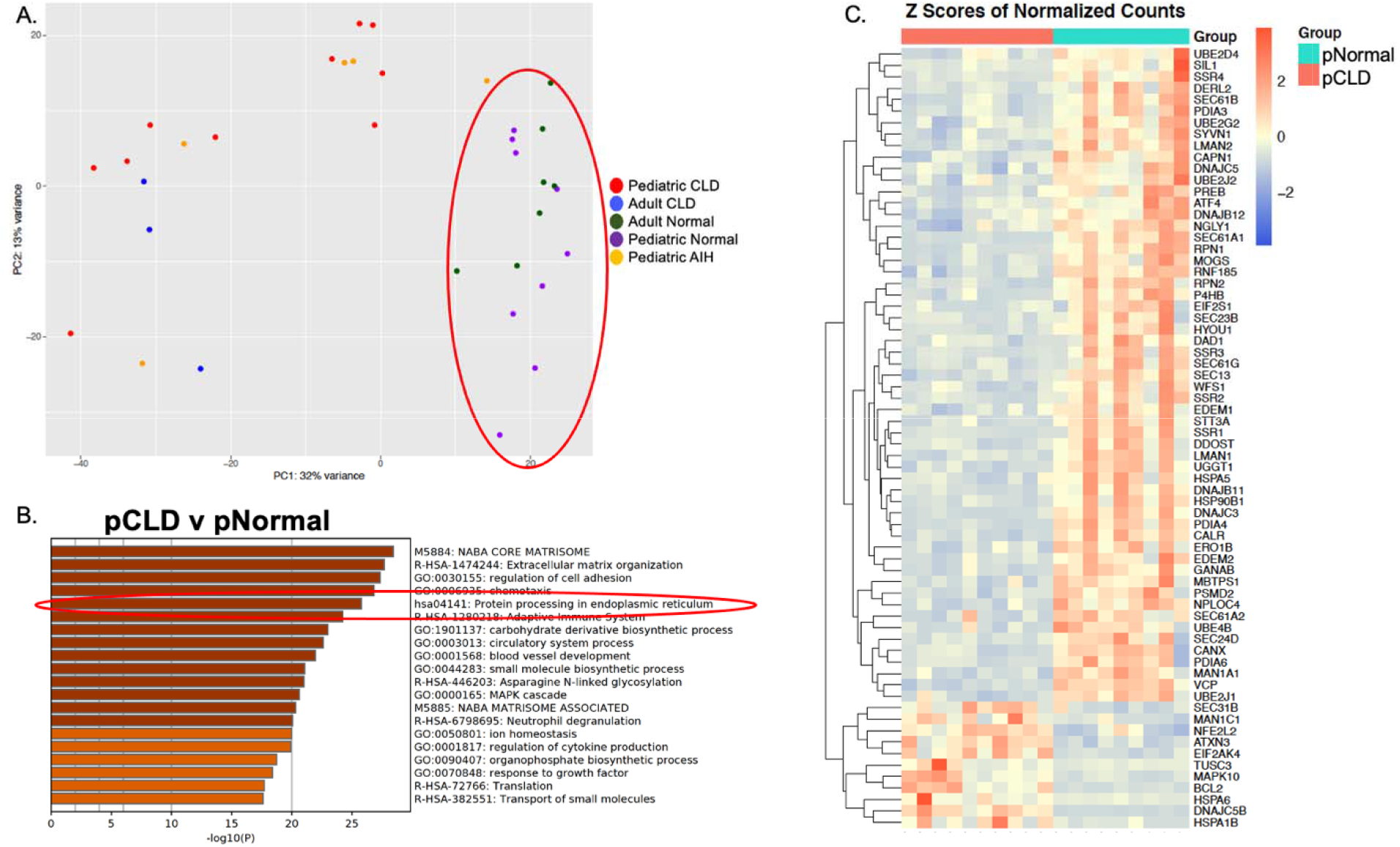
Principal Component Analysis (PCA) and Metascape pathway analysis of differentially expressed genes in pediatric cholestatic liver disease (pCLD) samples (n=10) compared to pediatric normal (pNormal) samples (n=9). A) PCA of RNA-seq data demonstrated that all pediatric and adult disease samples (CLD and AIH) cluster distinctly from normal pediatric and normal adult controls (circled in red). B) Metascape pathway analysis of RNA-seq differentially expressed genes comparing pCLD to pNormal samples demonstrated that genes involving the KEGG pathway hsa04141 “*protein processing in endoplasmic reticulum*” (circled in red) was among the top 5 differentially expressed pathways (−log10(P)=25.77). C) Heatmap demonstrating relative expression of differentially expressed genes within KEGG pathway hsa04141 AIH=autoimmune hepatitis

### Hepatic IRE1α/XBP1 pathway gene and protein expression is decreased in pCLD compared to pNormal livers

Since cholestasis and elevated levels of bile acids can induce hepatic ER stress, we sought to identify if there were differences in the expression of specific UPR genes and proteins between pCLD and pNormal livers. Quantitative PCR analysis of IRE1α/XBP1 pathway genes demonstrated that the pCLD group had a 43% decrease in *IRE1α* expression (1.4 ± 0.2 vs 0.8 ± 0.2, p<0.05), and a 67% decrease in *XBP1s* expression (1.5 ± 0.3 vs 0.5 ± 0.1, p<0.01), compared to pNormal livers (Fig. 2A). Additionally, gene expression of the XBP1s downstream targets endoplasmic reticulum DNA J domaincontaining protein 4 (*ERdj4*) and ER degradation enhancing α-mannoside 1 (*EDEM1*) were also lower in pCLD samples compared to pNormal livers (Fig. 2A). *EDEM1* was decreased by 73% (1.5 ± 0.2 vs 0.4 ± 0.07, p<0.001) and *ERdj4* was decreased by 50% (1.0 ± 0.2 vs 0.5 ± 0.07, p<0.05). Gene expression of liver *IRE1α, XBP1s, EDEM1* and *ERdj4* were similar in PFIC and ALGS, although there was a trend toward lower gene expression in the PFIC samples (Fig. 2B).

**Figure 2.**
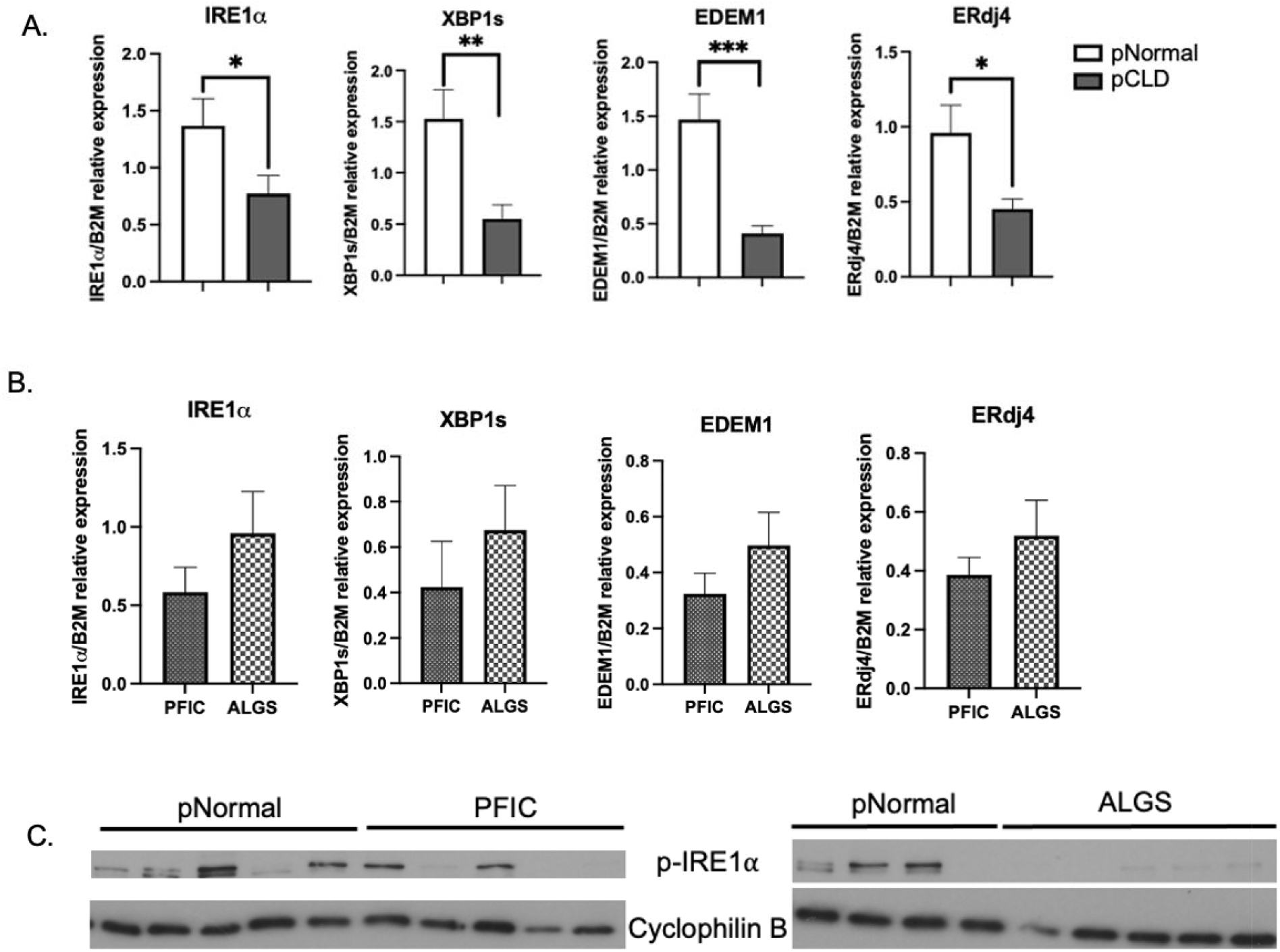
Hepatic IRE1α/XBP1 pathway of the UPR is decreased in pediatric cholestatic liver disease (pCLD) samples compared to pediatric normal (pNormal) livers. A) Hepatic gene expression of IRE1α, XBP1s and downstream targets EDEM1 and ERdj4 were measured via qPCR. pCLD livers (n=10) had decreases in IRE1α, XBP1, EDEM1 and ERdj4 expression compared to pNormal livers (n=9), (*p<0.05), **p<0.01, ***p<0.001) B) A sub-group analysis of pCLD samples demonstrated no change in hepatic gene expression of IRE1α, XBP1s, EDEM1 and ERdj4 via qPCR when comparing Alagille (ALGS) (n=5) to progressive familial intrahepatic cholestasis (PFIC) cohorts (n=5). C) Protein expression of p-IRE1α was decreased in pCLD (both pediatric PFIC and ALGS) patients compared to pNormal livers (n=9).

Analysis of protein expression demonstrated decreased expression of phosphorylated IRE1α (p-IRE1α) in pCLD (in both ALGS and PFIC) samples compared to pNormal controls (Fig. 2C) while phosphorylated PERK (p-PERK) could not be detected. The p-PERK target phosphorylated eukaryotic translation initiation factor 2α (p-eIF2α) and downstream activating transcription factor 4 (ATF4) had higher protein expression in the pCLD (in both ALGS and PFIC) group, compared to pNormal controls, suggestive of ongoing ER stress (Fig. 3A, B). Finally, ATF6 gene expression in pCLD and pNormal controls was similar, with trends toward lower ATF6 expression in the pCLD group (p=0.09) (Fig. 3C) and a similar trend when pNormal was compared to either ALGS or PFIC sub-groups (Supplemental Fig. 4B).

**Figure 3.**
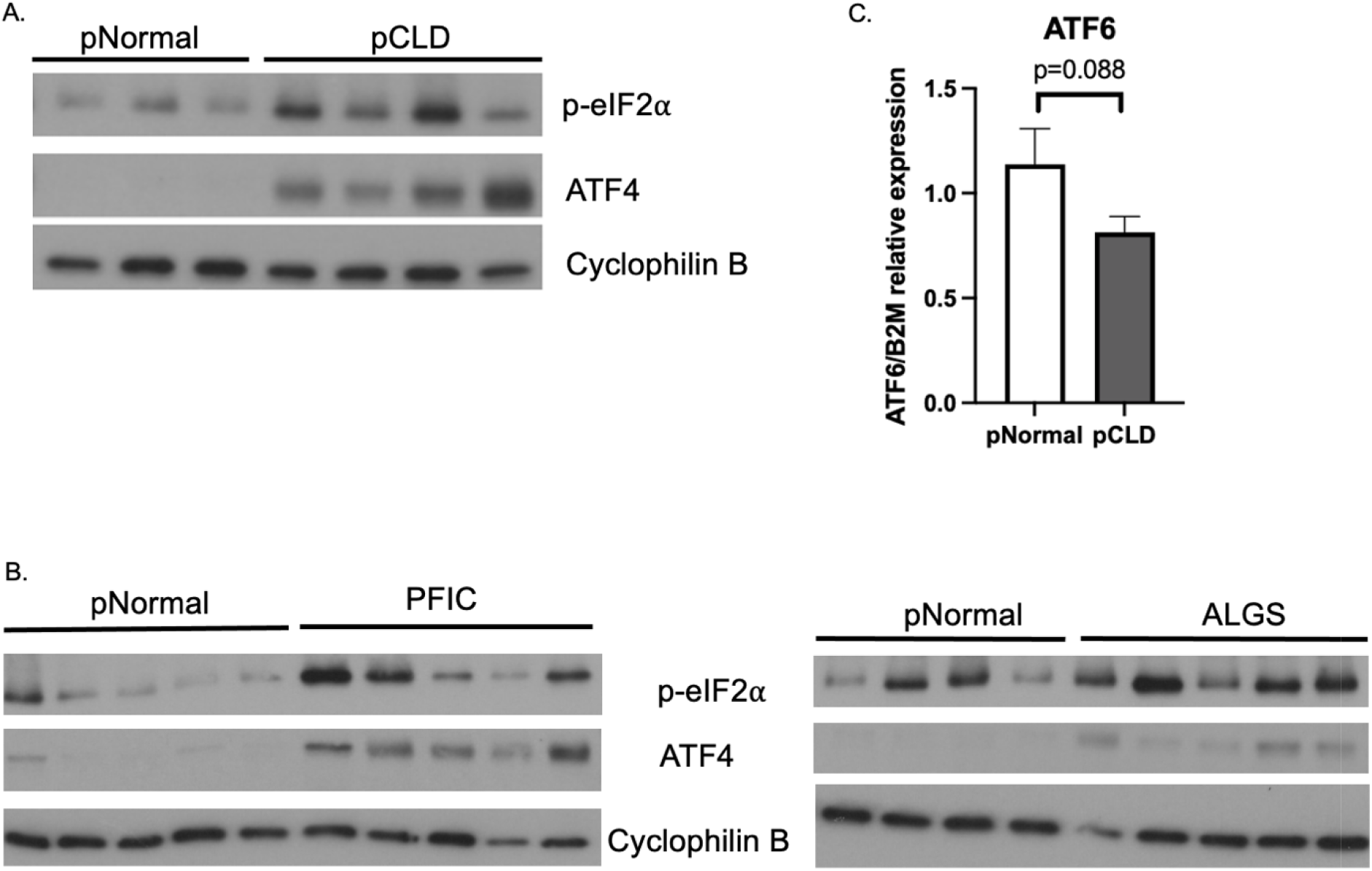
Hepatic protein expression of PERK pathway targets and gene expression of ATF6 in pediatric cholestatic liver disease (pCLD) livers compared to pediatric normal (pNormal) controls. A) Western blotting demonstrated an increase in protein expression of phosphorylated eIF2α (p-eIF2α) and ATF4 in pCLD livers (pooled n=2-3 samples per lane) compared to pNormal livers (pooled n=2-3 samples per lane). B) Increased p-eIF2α and ATF4 protein expression in Alagille (ALGS) and progressive familial intrahepatic cholestasis (PFIC) samples compared to pNormal livers (n=9 total, split between 2 blots). C) qPCR demonstrated a trend towards decreased ATF6 gene expression in pCLD livers (n=10) compared to pNormal livers (n=8), but this did not reach statistical significance (1.1 ± 0.2 vs 0.8 ± 0.08, p=0.088).

### Hepatic pathway analysis of RNA-seq from patients with AIH compared to pCLD patients and pNormal controls

In order to evaluate which UPR pathway findings in our pCLD were cholestatic liver disease-specific and/or related to progressive liver disease, we utilized explanted livers from patients with AIH as a disease control for advanced, fibrotic non-cholestatic liver disease. RNA-seq data comparing pCLD and AIH disease controls identified 3643 differentially expressed genes. Pathway analysis of the top 3000 differentially expressed genes did not demonstrate any differences related to protein processing in the ER pathways, ER stress or other UPR-related pathways (Fig. 4A). In contrast, RNA-seq comparing pediatric AIH and pNormal control samples identified 3608 differentially expressed genes and pathway analysis of differentially expressed genes between AIH and pNormal groups identified that KEGG pathway hsa04141 “protein processing in endoplasmic reticulum” was among the top 10 differentially expressed pathways (−log10(P)=23.33) (Fig. 4B), the same pathway detected when pCLD and pNormal were compared. Figure 4C is a heatmap demonstrating that the majority of the genes within this pathway were decreased in the AIH samples compared to pNormal. Metascape pathway analysis of all downregulated or all upregulated genes in AIH compared to pNormal samples (n=1579 and n=2029 respectively) identified that KEGG pathway hsa04141 was significantly enriched in down-regulated genes, confirming that this pathway is downregulated in AIH samples (Supp. Fig. 5A and B).

**Figure 4.**
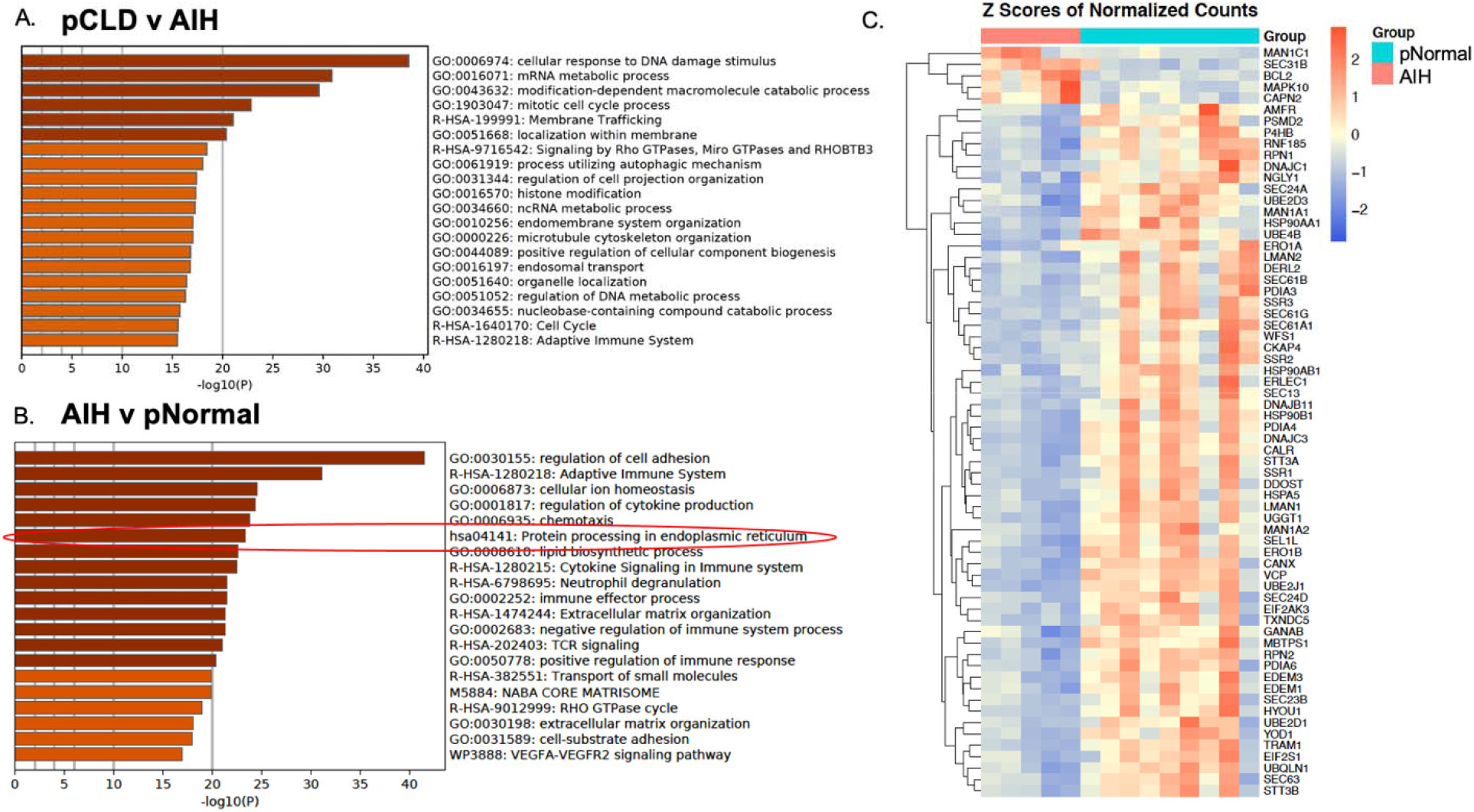
Pathway analysis of pediatric autoimmune hepatitis (AIH) samples (n=5) compared to pediatric normal (pNormal) controls (n=9). A) Metascape pathway analysis of RNA-seq differentially expressed genes comparing pCLD to AIH livers showing the top 20 differentially expressed pathways. B) Metascape pathway analysis of RNA-seq differentially expressed genes comparing AIH to pNormal livers demonstrated that the KEGG pathway hsa04141 “*protein processing in endoplasmic reticulum*” (circled in red) was significantly differentially expressed (−log10(P)=23.33). C) Heatmap demonstrating expression of differentially expressed genes within KEGG pathway hsa04141 between AIH and pNormal livers.

### Hepatic gene and protein expression from patients with AIH compared to pCLD patients and pNormal controls

We next evaluated whether gene and protein expression of canonical UPR pathways differed between the pCLD and AIH cohorts. When compared to the AIH group, pCLD livers had significantly lower gene expression of the IRE1α/XBP1 pathway genes *IRE1α, XBP1s, and EDEM1* by 78%, 77% and 67% (p<0.001, p<0.01, p<0.001, respectively) (Fig. 5A), although *ERdj4* gene expression was similar (Fig. 5A). Hepatic gene expression of ATF6 was also decreased by 43% in pCLD livers compared to AIH livers (0.8 ± 0.08 vs 1.4 ± 0.2, p<0.01) (Fig. 5B), while there were no differences in gene or protein expression of PERK, p-eIF2α or ATF4 (Fig. 5C, D).

**Figure 5.**
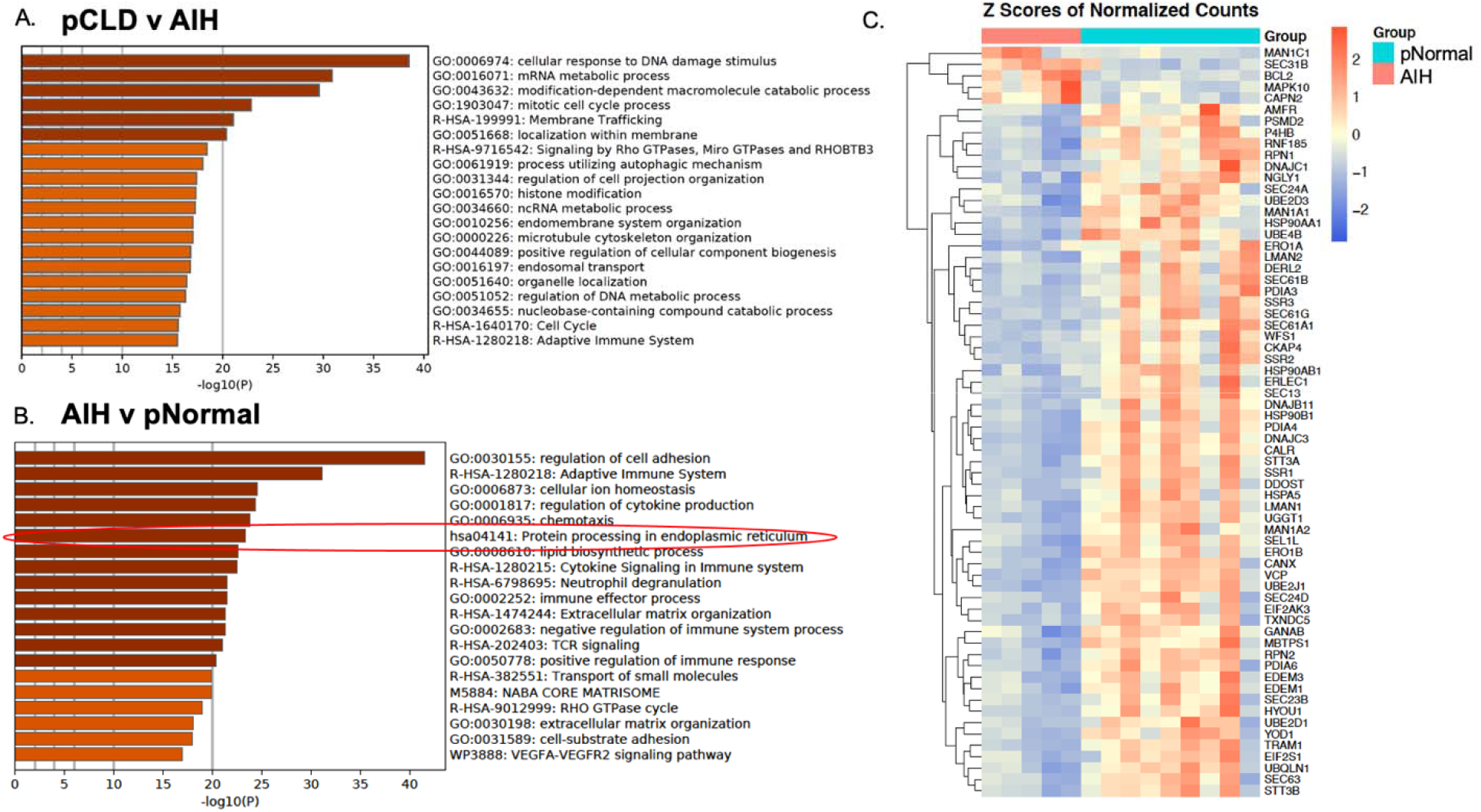
UPR gene and protein expression of autoimmune hepatitis (AIH) livers compared to pediatric normal controls (pNormal) and pediatric cholestatic liver disease (pCLD) samples. Using qPCR and Western blotting, we determine that: compared to AIH livers (n=5), pCLD livers (n=10) had decreased expression of IRE1α, XBP1s, and EDEM1 by 78%, 77%, and 67% (**p<0.01, ***p<0.001) while ERdj4 gene expression was similar. A) Gene expression of ATF6 was significantly decreased in pCLD livers compared to AIH livers (). B) There was no change in gene expression of PERK between pCLD and AIH livers. C) There was no significant change in protein expression of p-eIF2α and ATF4 in pCLD compared to AIH livers (n=2-3 samples pooled per lane).

Given these changes in the IRE1α/XBP1 pathway, we next compared AIH livers to our normal control samples. Gene expression of *XBP1s*, and *EDEM1* were similar between pNormal and AIH livers, while *IRE1α* expression was actually increased in the AIH group (3.5 ± 0.8 vs 1.4 ± 0.2, p<0.01) (Supp. Fig. 6A). ERdj4 expression was decreased in AIH samples compared to pNormal samples (0.3 ± 0.1 vs 1.0 ± 0.2, p<0.05) (Supp. Fig. 6A) similar to the pCLD findings. Protein expression of p-eIF2α and ATF4 was increased in AIH samples compared to controls suggestive of ongoing ER stress, while there was no difference in ATF6 gene expression between AIH and pediatric normal controls (Supp. Fig. 6B and 6C).

### Pathway analysis and hepatic UPR expression in aCLD liver explants

We next sought to evaluate if the changes identified in the pediatric CLD livers occurred in the adult CLD cohort. There were 4,865 differentially expressed genes in aCLD compared to aNormal livers. Metascape pathway analysis again identified the KEGG pathway hsa04141 in the top 100 differentially expressed pathways (−log10(P)=11) (Fig. 6A) and identified this as a downregulated pathway (Supp. Fig. 7A and 7B). This pathway was not differentially expressed between pCLD and aCLD cohorts (Fig. 6B). Although IRE1α/XBP1 pathway gene expression in aCLD compared to aNormal showed a trend toward decreased gene expression of *IRE1α, XBP1s, ERdj4* and *EDEM1*, these differences were not statistically significant (Fig. 6C). Gene expression of *ATF6* between aCLD and normal samples was similar (Fig. 6C), and there was an increase in protein expression of p-eIF2α and ATF4 in the aCLD samples compared to controls, similar to the pCLD experiments (Fig. 6D).

**Figure 6.**
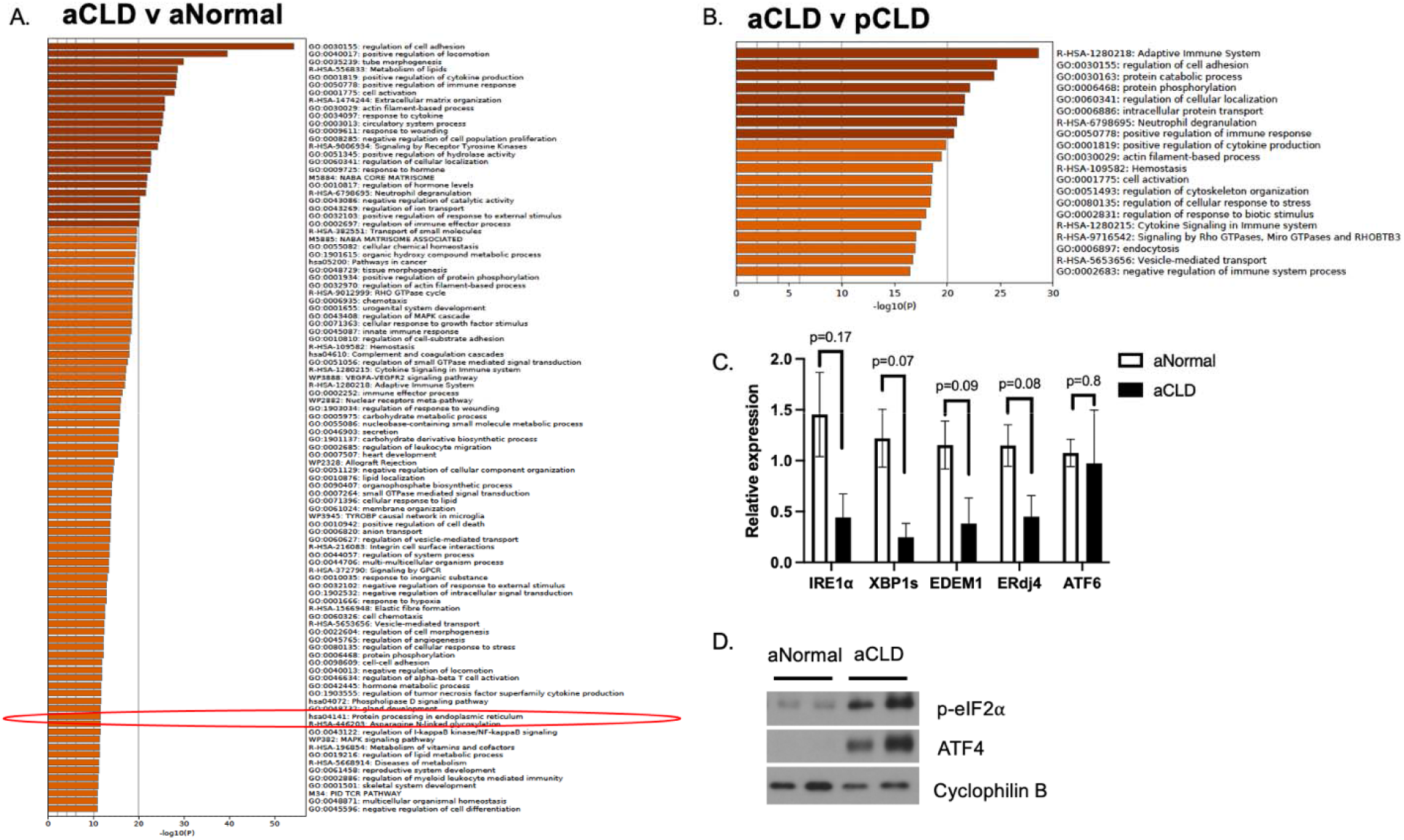
Pathway analysis and UPR expression of adult cholestatic liver disease (aCLD) livers compared to adult normal controls (aNormal). A) Metascape pathway analysis of RNA-seq differentially expressed genes comparing aCLD (n=3) to aNormal (n=7) samples demonstrated that the KEGG pathway hsa04141 “protein processing in endoplasmic reticulum” (circled in red) was among the top 100 differentially expressed pathways(−log10(P)=11). B) Metascape pathway analysis of RNA-seq differentially expressed genes comparing pCLD (n=10) to aCLD (n=3) samples. The top 20 differentially expressed pathways are shown. C) qPCR demonstrated that hepatic gene expression of IRE1α, XBP1s, EDEM1 and ERdj4 in aCLD livers were similar to pNormal controls. There was no change in hepatic gene expression of ATF6 between aCLD and normal controls. D) There was an increase in protein expression of p-eIF2α and ATF4 in aCLD samples (3 samples, n=1-2 per lane) compared to aNormal livers (pooled samples, n=3 per lane).

## Conclusion

Pediatric cholestatic liver diseases are the leading indication for pediatric liver transplantation and there are currently no effective medical therapies that reduce the need for liver transplantation. During cholestasis, high intrahepatic concentrations of bile acids increase ER stress and hepatocellular injury, and an effective UPR response is essential to restore cellular homeostasis ^2–5^ Inadequate activation or dysregulation of the IREα/XBP1 pathway of the UPR has been implicated in the pathogenesis of several liver diseases^23^. Puri et al demonstrated that decreased levels of XBP1s protein expression was associated with the development of steatohepatitis in adult patients, although, although there remains some conflicting data^8, 24, 25^. In addition, the IRE1α/XBP1 pathway has been implicated in in murine models of pharmacologic or fatty liver-induced ER stress^14, 26^. Henkel et al demonstrated that following tunicamycin-induced ER stress, liver specific XBP1 knock out (LS-XBP1^-/-^) and XBP1^fl/fl^ mice have similar initial activation of the UPR, but LS-XBP1^-/-^ mice have impaired resolution of ER stress with resultant apoptosis^14^. Recently, our group utilized a bile acid feeding model of cholestasis and bile acid toxicity to demonstrate that, compared to adult mice, young mice had decreased XBP1s downstream target activation, with a resultant enhanced susceptibility to bile acid induced liver injury and apoptosis^15^. While select UPR genes can identify adult PSC patients who are at high risk for liver-related complications, there is no other data on the role of the UPR in human cholestatic liver disease nor in pediatric cholestatic liver diseases^27^. Therefore, we sought to determine the level of UPR activation in a pediatric cholestatic liver disease cohort. Based on our murine findings in young mice, we hypothesized that pediatric cholestatic livers would have lower levels of IRE1α/XBP1 pathway expression compared to normal control or disease-control livers.

Unbiased gene expression pathway analysis demonstrated the novel finding that the “protein processing in endoplasmic reticulum” KEGG pathway was decreased in pediatric cholestatic livers compared to normal controls. In addition, 4 other highly significant pathways included ‘NABA CORE MATRISOME’, the reactome pathway: ‘Extracellular matrix organization’, and the GO Biological Processes pathways: regulation of cell adhesion’ and ‘chemotaxis’. These pathways include genes primarily related to extracellular matrix organization including collagens and proteoglycans, as well as genes related to leukocyte migration and cytokines. Given the fibroinflammatory nature of ALGS and PFIC diseases, these pathway changes are not surprising. However, the finding of a pathway related to protein processing within pediatric cholestatic liver diseases has not previously been described.

We next demonstrated that hepatic IRE1α protein expression and IRE1α/XBP1 pathway downstream target genes were decreased in pediatric cholestatic disease livers compared to normal controls. This impaired or dysregulated hepatic IRE1α/XBP1 pathway expression is consistent with previously reported findings in murine models of cholestasis and murine and human fatty liver disease ^12, 15, 24, 26^. Specifically, ERdj4 and EDEM1 are essential for the process of ER associated degradation (ERAD) where misfolded proteins are targeted for degradation, and ERdj4 deficiency in cells can lead to constitutive ER stress^28–30^. Our pediatric cholestatic disease cohort included patients with ALGS and PFIC. Although our sample size was limited, we did not find any differences between our ALGS and PFIC samples related to ER stress or UPR pathway signaling. This indicates that one pediatric cholestatic liver disease was not primarily responsible for our findings and suggests that the observed findings are due to cholestatic liver disease, rather than specific disease etiology.

In order to determine whether the IRE1α/XBP1 pathway findings in pCLD explant samples were due to the effects of cholestasis and/or due to the advanced liver disease with extensive fibrosis, we next utilized explanted, cirrhotic livers from pediatric patients with AIH. In fact, despite having end-stage liver disease, 4 of 5 AIH patients had serum bilirubin levels of 2mg/dl or lower. We determined that expression of the IRE1α/XBP1s pathway was lower in pediatric cholestatic liver samples than levels in autoimmune hepatitis disease-control livers, indicating that these changes are likely due to the cholestatic liver diseases, rather than cirrhosis.

In contrast, some UPR changes in the pediatric cholestatic liver disease group identified using comparisons to normal livers were also noted when pCLD and AIH cohorts were compared, indicating that these effects are likely due to the presence of cirrhosis and end-stage liver disease. Both pCLD and AIH samples had evidence of increased p-eIF2α expression, which can lead to decreased protein translation, while selectively allowing translation of certain mRNAs including the transcription factor ATF4^31^. ATF4 promotes resolution of ER stress by activating target genes involved in antioxidant responses and amino acid synthesis, but also activates CCAAT-enhancer-binding protein homologous protein (CHOP), which can precipitate ER-stress mediated apoptosis^32^. Unfortunately, we were unable to detect CHOP expression in any of our liver samples. These findings are supportive of ongoing cellular or ER stress within the pCLD and AIH cohorts where eIF2α/ATF4 activation may be indicative of unresolved ER stress. This ER stress may be due, at least in part, to inadequate or dysregulated activation of the IRE1α/XBP1 pathway, with a resultant inability to adequately resolve ER stress as has been demonstrated in murine models^14, 15, 26^. Finally, there was a trend towards decreased ATF6 expression in the pCLD cohort compared to normal samples, though we were unable to detect ATF6 protein expression in our explanted livers.

It is unclear if the reduced hepatic IRE1α/XBP1 pathway expression seen in the cholestatic liver disease group is caused by the cholestasis and elicited only in end-stage liver disease, or whether these patients had an underlying genetic or developmental impairment of hepatic UPR activation which contributed to their progression to end-stage liver disease. Given the spectrum of clinical manifestations and disease progression seen within ALGS and PFIC patients that cannot be attributed solely to the underlying primary gene defects, our data is consistent with additional defects of IRE1α/XBP1 signaling which may impair ER stress resolution and increase these patients’ susceptibility to develop more rapid progression or more severe liver disease.

There are several potential limitations to this study. Adult cholestatic liver disease samples had similar trends of the IRE1α/XBP1 and p-eIf2a-ATF4 expression suggesting that the UPR changes in the cholestatic liver groups may not be restricted to pediatric populations. However, the adult sample sizes were small and therefore we cannot exclude the possibility that there were age-related differences in pediatric and adult cholestatic liver diseases that we were unable to identify. The ages of our pediatric CLD patients were not available, although LCH does not routinely perform liver transplants on patients above 18 years of age. Additionally, the liver explant samples were obtained from a biorepository which contained a very limited amount of clinical data and no post-transplantation patient outcomes. Since the study was performed in liver explants and analysis is therefore limited only one time point late in the progressive disease, we cannot determine whether the impaired XBP1 pathway activation was causative for disease progression, or potentially the consequence of it. Nonetheless, several UPR changes observed in livers with cholestatic liver disease differed from our AIH non-cholestatic disease control group, suggesting that they are more specific to the cholestatic nature of these diseases.

This study identified that hepatic gene and protein expression of IRE1α/XBP1 signaling and other UPR signaling differs in progressive cholestatic liver diseases and therefore may be important in the pathogenesis of disease. Although some changes in the UPR are related to cirrhosis, several are unique to patients with cholestatic liver disease. Since hepatic IRE1α/XBP1 pathway expression is protective in murine models of cholestatic liver injury, the attenuated expression of this canonical UPR pathway may be an important factor in disease progression. These data also identify UPR proteins and genes that may serve as putative targets for the development of novel therapeutic interventions.

