## Supplemental tables and figures for "IRE1α/XBP1 pathway expression is impaired in pediatric cholestatic liver disease explants"

Supplemental Table 1. Antibody List

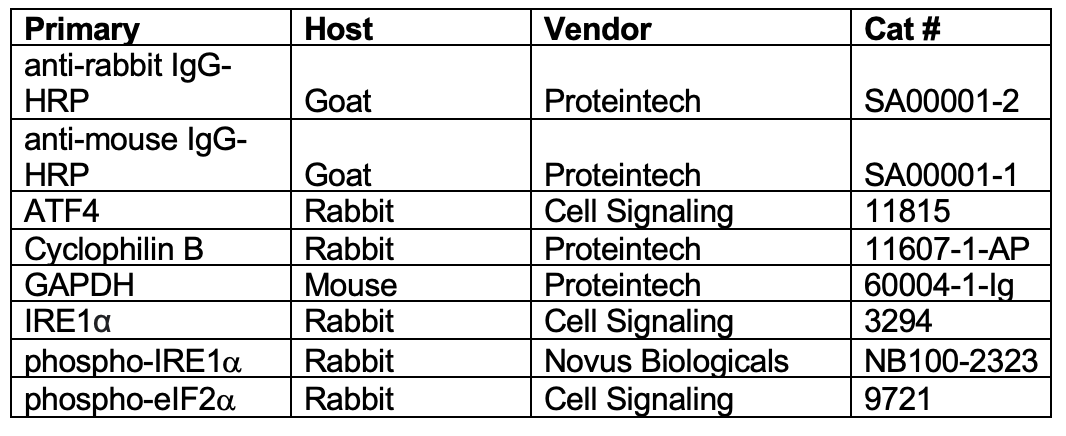

Supplemental Table 2. Primer sequences used in qPCR.

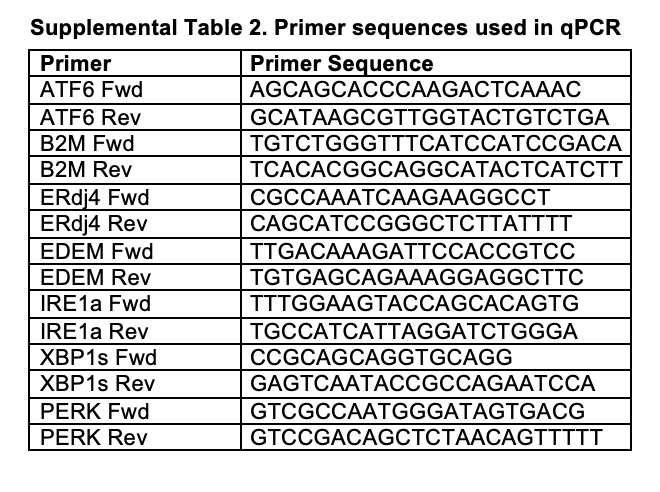

Supplemental Figure 1. Patient demographic and clinical data.
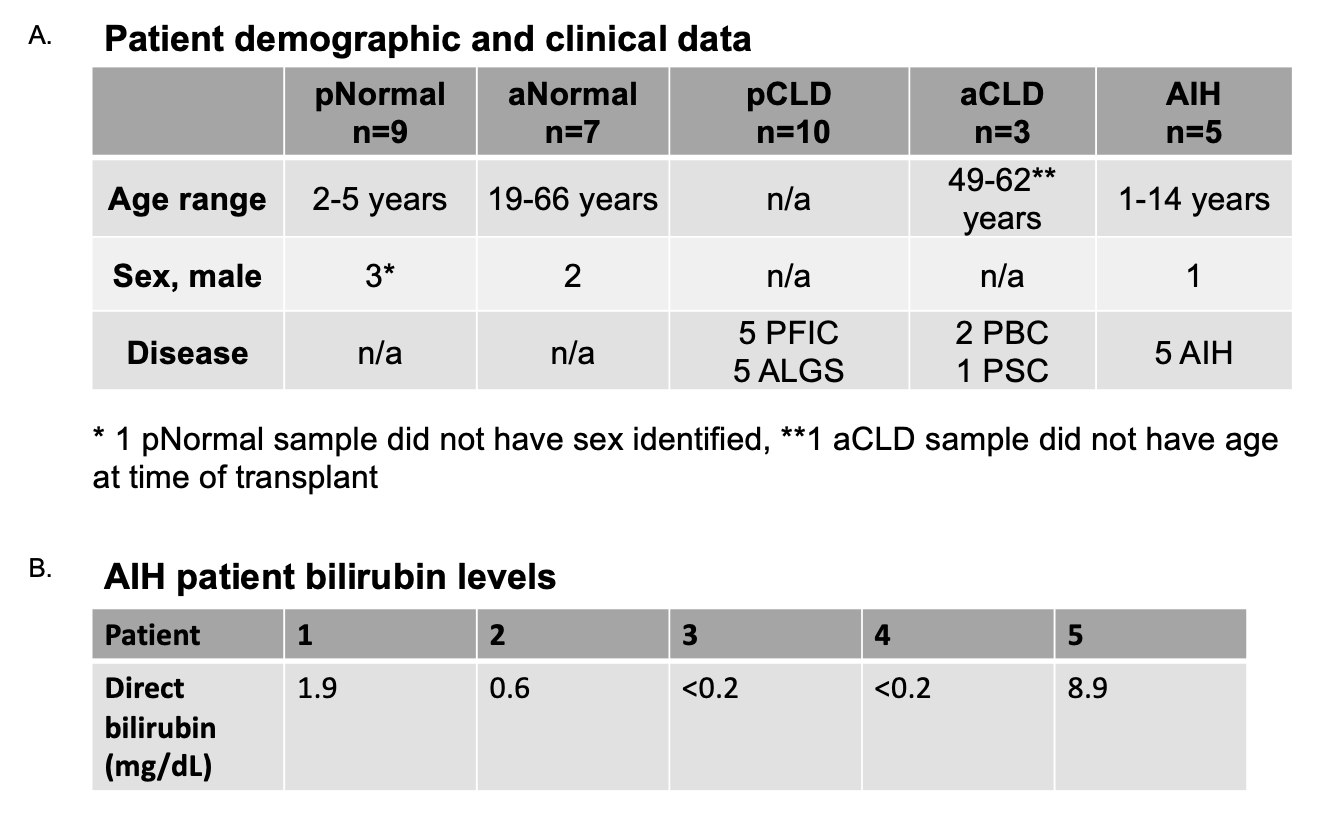

A) Demographic data of 34 human liver explant samples.
B) Direct bilirubin level (mg/dL) of autoimmune hepatitis (AIH) patients (n=5) on the day prior to liver transplantation.

pNormal = pediatric normal liver, aNormal = adult normal liver, pCLD = pediatric cholestatic liver disease, aCLD = adult cholestatic liver disease, AIH = autoimmune hepatitis, mos = months

Supplemental Figure 2. METAVIR fibrosis scores.
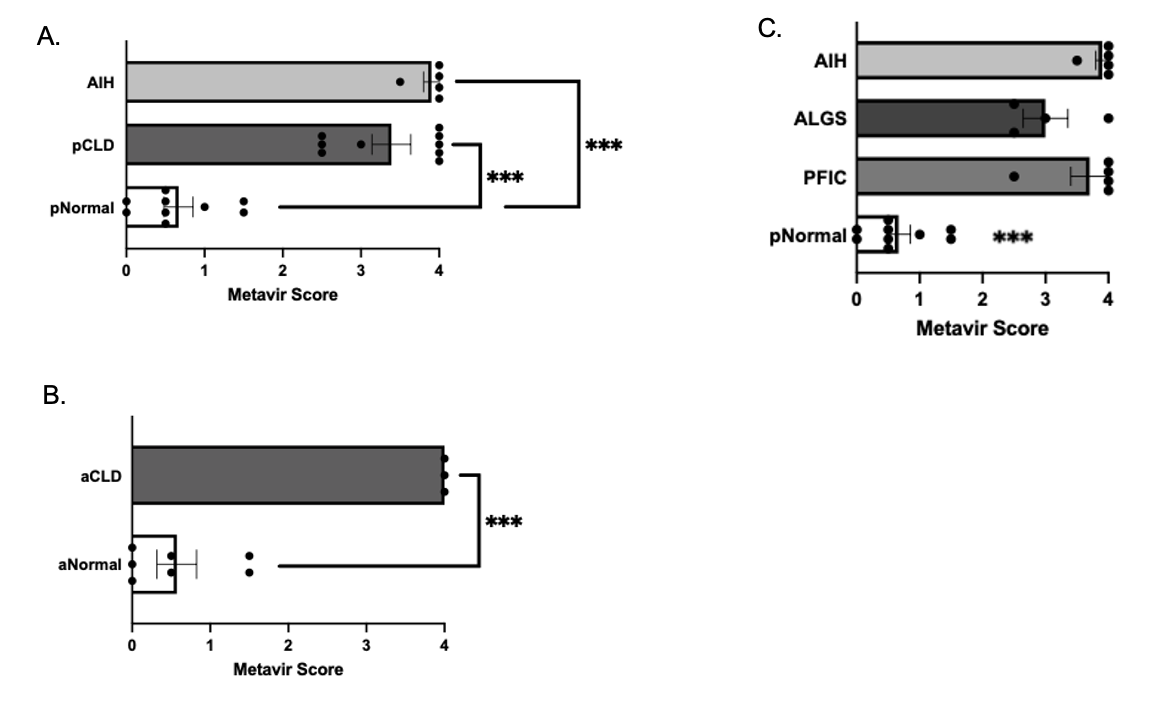

A) Pediatric cholestatic liver disease (pCLD) (n=9) and autoimmune hepatitis (AIH) livers (n=4) demonstrated significantly more fibrosis than pediatric normal samples(pNormal) (n=9) (***p<0.001).
B) Adult cholestatic liver disease (aCLD) livers (n=3) had significantly more fibrosis than adult normal (aNormal) samples (n=7) (***p<0.001).
C) Alagille (ALGS) (n=4) and progressive familial intrahepatic cholestasis (PFIC) samples (n=5) had significantly more fibrosis than pNormal (n=9) and had similar METAVIR scores to AIH samples (n=4) (***p<0.001).

Supplemental Figure 3.

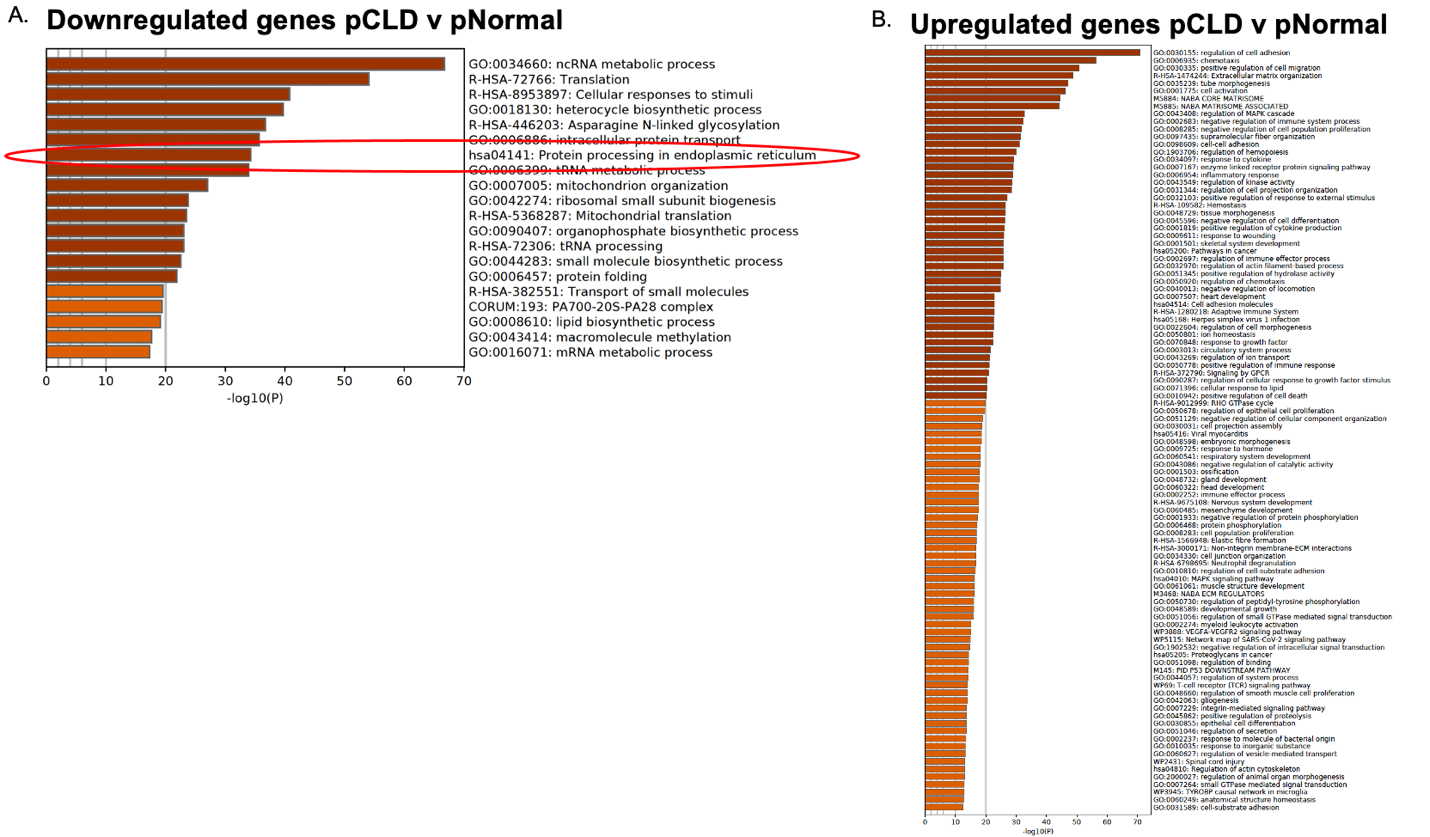

Supplemental Figure 3. Metascape pathway analysis of significant downregulated and upregulated genes in pediatric cholestatic liver disease (pCLD) samples (n=10) compared to pediatric normal (pNormal) samples (n=9).

A) Utilizing all significantly different downregulated genes in pCLD for pathway analysis, the KEGG pathway hsa04141 *“protein processing in endoplasmic reticulum”* (circled in red) was among the top 10 differentially expressed pathways (-log10(P)=34.28).
B) Utilizing the top significantly upregulated genes in pCLD for pathway analysis KEGG pathway hsa04141 *“protein processing in endoplasmic reticulum”* was not differentially expressed.

Supplemental Figure 4.

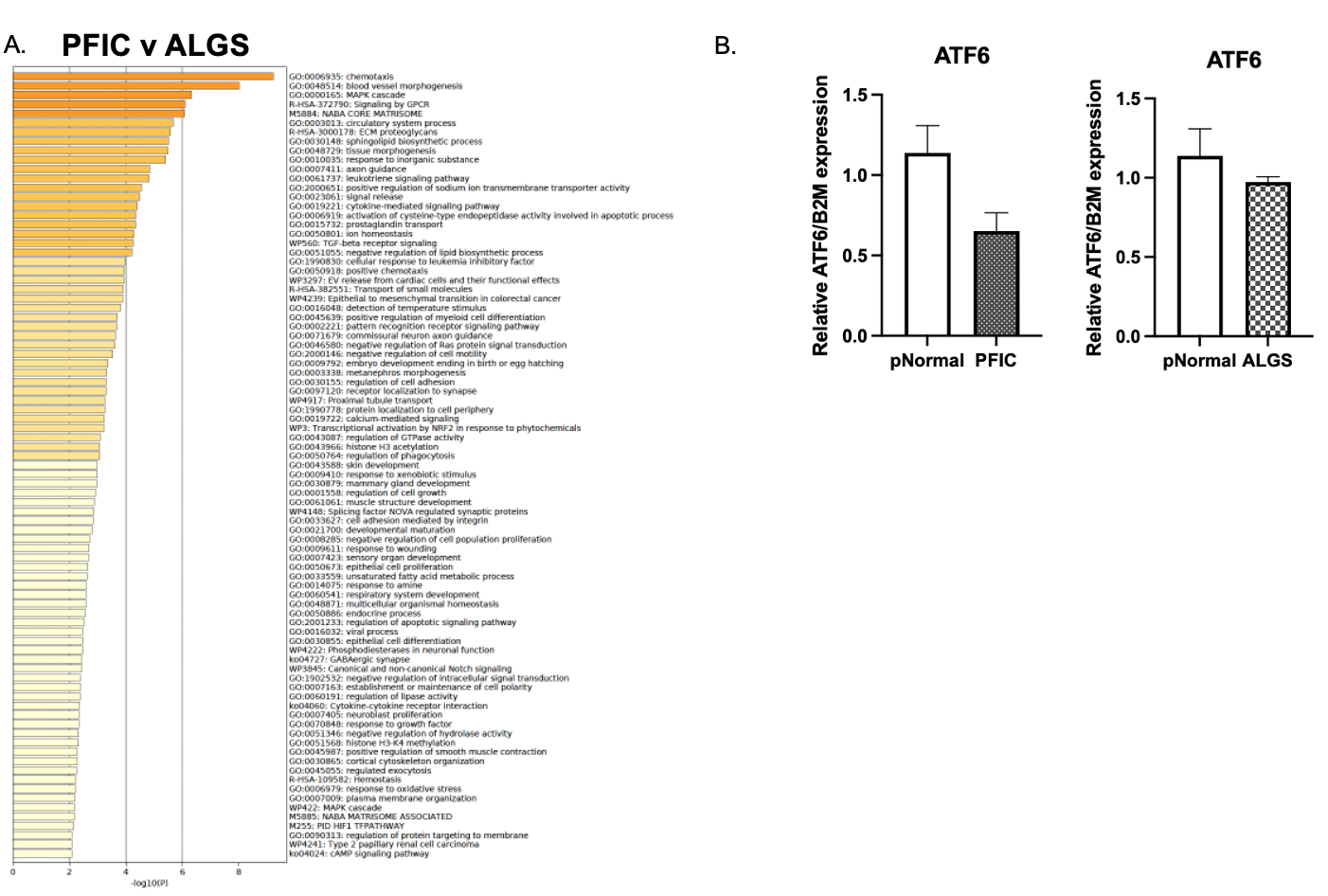

Supplemental Figure 4. Metascape pathway analysis on differentially expressed genes between progressive familial intrahepatic cholestasis (PFIC) and Alagille (ALGS) samples.

A) The top 100 pathways enriched in differentially expressed genes.
B) There was no decreased ATF6 gene expression in PFIC (n=5) and ALGS (n=5) livers compared to pNormal livers (n=9) (p=0.07 and p=0.5 respectively).

Supplemental Figure 5.

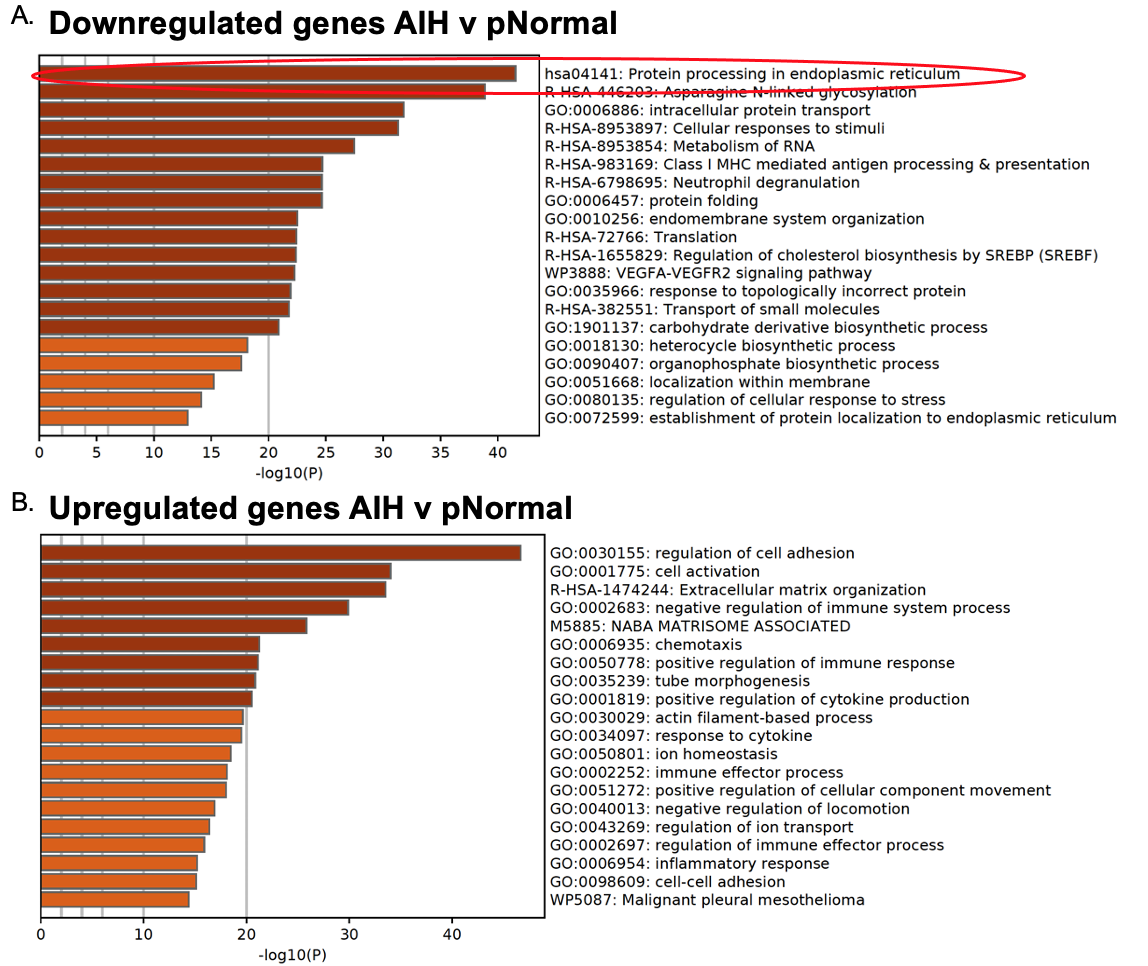

Supplemental Figure 5. Metascape pathway analysis of significant upregulated and downregulated genes in pediatric autoimmune hepatitis (AIH) samples (n=5) compared to pediatric normal (pNormal) samples (n=9).

A) Utilizing all significantly different downregulated genes in AIH samples for Metascape pathway analysis, the KEGG pathway hsa04141 *“protein processing it endoplasmic reticulum”* (circled in red) was the top differentially expressed pathway (-log10(P)=41.54).
B) Utilizing all significantly different upregulated genes in AIH samples for pathway analysis. The top 20 differentially expressed pathways are shown.

Supplemental Figure 6.
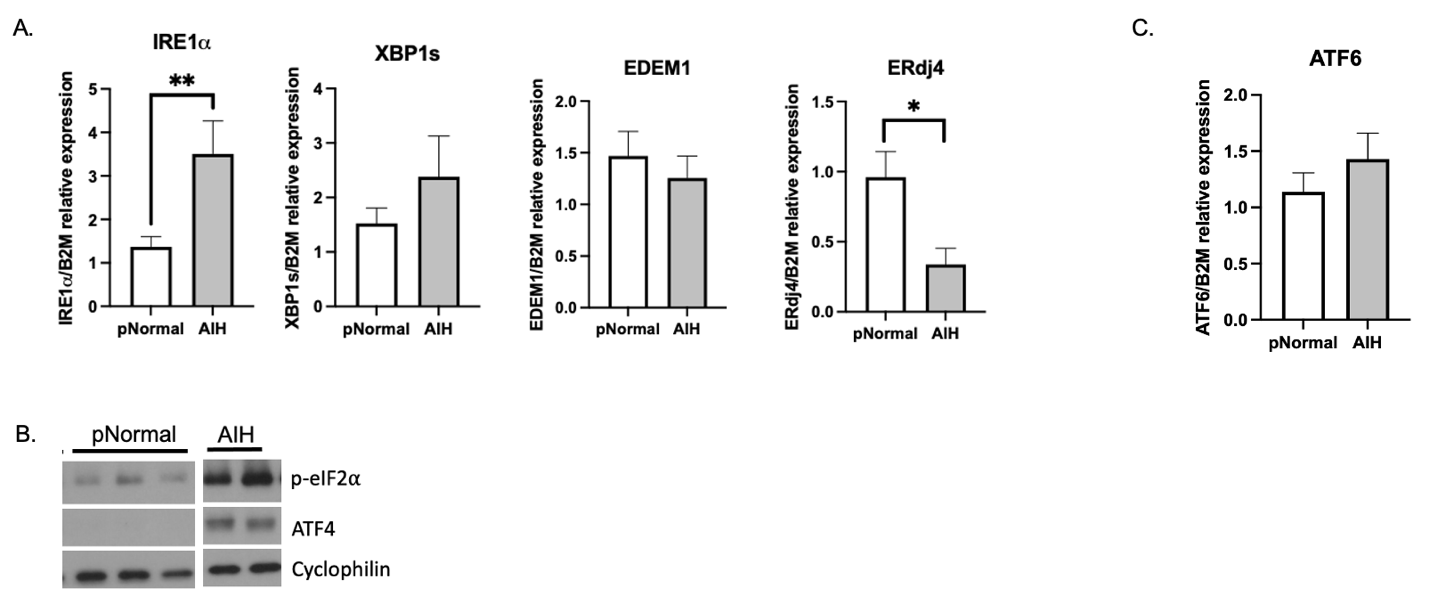

Supplemental Figure 6. UPR gene and protein expression of autoimmune hepatitis (AIH) samples compared to pediatric normal controls (pNormal).

A) qPCR demonstrated no difference in gene expression of XBP1s, and EDEM1 between pNormal (n=9) and AIH (n=5) livers, and in AIH livers, there was an increase in IRE1α expression and a decrease in ERdj4 expression compared to pNormal livers (*p<0.05 **p<0.01).
B) Western blotting demonstrated an increase in protein expression of phosphorylated p-eIF2α and ATF4 in AIH livers (pooled n=2-3 samples per lane) compared to pNormal livers (pooled n=2-3 samples per lane).
C) qPCR demonstrated that hepatic ATF6 gene expression was similar in AIH (n=5) and pNormal livers (n=9).

Supplemental Figure 7.
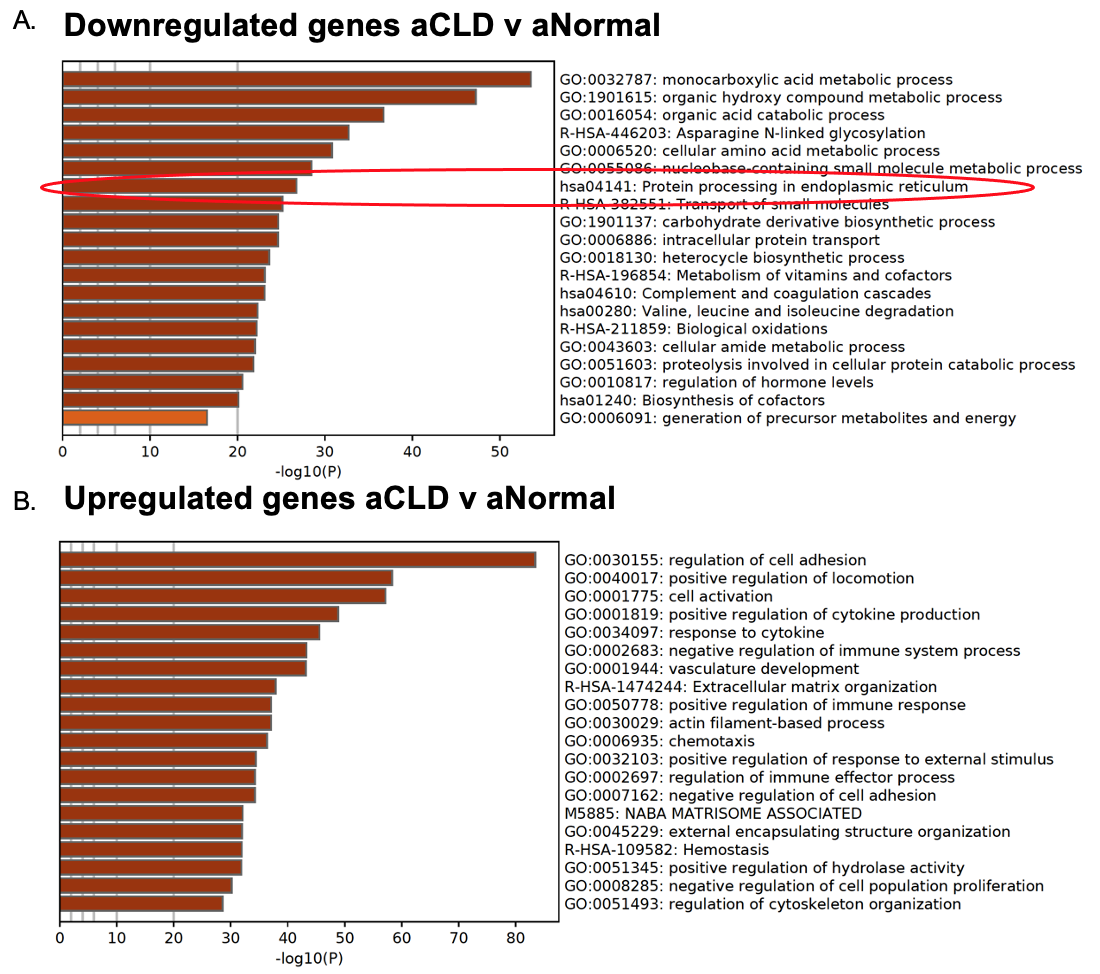

Supplemental Figure 7. Metascape pathway analysis of significant upregulated or downregulated genes in adult cholestatic liver disease (aCLD) samples (n=3) compared to adult normal (aNormal) samples (n=7).

A) Utilizing all significantly different downregulated genes in aCLD samples for Metascape pathway analysis, the KEGG pathway hsa04141 *“protein processing it endoplasmic reticulum”* (circled in red) was the top 10 differentially expressed pathway (-log10(P)=26.74).
B) Utilizing all significantly different upregulated genes in aCLD samples for Metascape pathway analysis, KEGG pathway hsa04141 *“protein processing in endoplasmic reticulum”* was not in the top 100 pathways enriched in these genes. The top 20 differentially expressed pathways are shown.
